# *VERNALIZATION2 (VRN2)* alters early tiller development in a facultative spring hexaploid bread wheat (*Triticum aestivum*)

**DOI:** 10.1101/2025.04.30.651433

**Authors:** Dominique Hirsz, Harry Taylor, India Lacey, Adam Gauley, Laura Dixon

## Abstract

Plants coordinate growth and development with environmental signals. Specifically, temperature and photoperiod are important regulators to time the transition from vegetative to floral growth. Through utilising these environmental signals potential reproductive success is optimised, which is critical for the production of high crop yields. For cereals, some require a winter before undergoing reproductive development. This extended period of cold exposure enables the process of vernalization and is also important for the synchronised timing of the floral transition. The cereal-specific floral repressor *VERNALIZATION2 (VRN2)* has an integral role in this pathway, yet this locus remains poorly characterised in hexaploid wheat. Our research in a facultative wheat suggests that the tandemly duplicated genes comprising the *VRN2* locus, *ZCCT1* and *ZCCT2*, respond to multiple environmental factors and have differences in gene expression patterns at a gene and sub-genome level. These genes also show co-regulation, forming a regulatory loop between *ZCCT-D1* and *ZCCT-D2*. The function of these genes beyond the classic vernalization response is explored in a facultative wheat with *VRN-D2* identified to regulate early tiller development, with an accelerated rate of secondary tiller emergence and presence of coleoptile tillers. This phenotype increases ground coverage more rapidly, which provides agronomic benefits including improved competition with biological stressors. The findings identify that the *VRN2* loci in bread wheat is formed of multiple genes which have overlapping but also unique regulation and function. Selecting these genes individually may offer a route to alter wheat plant architecture without directly impacting vernalization requirement.

**Highlight:** 

## Introduction

Bread wheat *(Triticum aestivum)* is the most widely cultivated cereal crop, made possible by allelic variations in several key genes which have been enriched for during selection and breeding. However, bread wheat remains extremely vulnerable to the effects of climate change. Each global temperature increase of 1°C is expected to result in global losses in yield of 6%, with anticipated losses ranging from approximately 4-20% for different regions [1, 2]. The ability to improve and modify the adaptability of wheat to changing climate conditions is vital to maintain yield potential and support a rapidly increasing global population and food demand [3].

One aspect of adaptability which impacts yield potential is the vernalization requirement. This is the requirement for a prolonged exposure to cold temperatures, coupled with short-day photoperiods, which provides plants with a means for monitoring winter and timing reproductive development to favourable conditions. Vernalization is quantitative, so the length of exposure to cold temperatures directly determines the rate of floral transition. Flowering time is accelerated following a longer cold duration up to a point, after which the vernalization requirement is satisfied and cold temperatures have limited or negative impact on flowering date [4]. In countries with a suitable climate pattern to meet the vernalization requirement, vernalization-requiring (termed ‘winter’ wheat) has a higher yield potential than other wheat [5, 6]. Furthermore, it provides a number of other agronomic benefits including establishment and ground cover over winter and competition against weeds such as blackgrass [7-9].

Wheat varieties which do not require vernalization are termed ‘spring’ varieties, avoiding the pathway through suppression of floral repressors or constitutive expression of floral activators [4, 10, 11]. A further group exists which are facultative wheats, these do not require vernalization to flower but flowering time is accelerated when they experience vernalization. Changing environmental patterns are altering previously established agricultural practices with the optimal timing of crop planting becoming less predictable. For winter crops the variable winter temperatures are making the timing for the completion of vernalization unpredictable. This presents two major challenges; too early completion causing a shift in meristem development and a reduction in plant biomass; or too late completion delaying flowering and in extreme cases not being satisfied before warmer spring conditions and so a substantial reduction in final yield. Increasingly, facultative wheats are being used which can balance these requirements.

Understanding the contributions of the genetic components of the vernalization pathway is important to enable the development of wheat with a variety of vernalization responses. The genetic core of the vernalization pathway in cereals involves *VERNALIZATION1 (VRN1), VERNALIZATION2 (VRN2)* and *FLOWERING LOCUS T1 (FT1)*, also called *VRN3*. Mutations in the promoter or first intron of *VRN1* lead to overexpression, which results in a dominant spring wheat phenotype [4]. *VRN2* is a monocot-specific floral repressor, highly expressed prior to vernalization and gradually repressed following cold exposure by the increasingly expressed *VRN1* [12]. Although *VRN2* is generally referred to as a gene, it is a locus made of two tandemly duplicated genes, *ZCCT1* and *ZCCT2* [13]. It has a putative C_2_H_2_ zinc finger domain and a CCT domain, named after the genes *CONSTANS, CONSTANS-like*, and *TIMING OF CAB 1* identified in *Arabidopsis thaliana*. Notably, *VRN2* is part of a known translocation between chromosome 4A and 5A [14]. Therefore, the *VRN2* gene in bread wheat is referring to a total of 6 genes, *ZCCT-A1* and *ZCCT-A2* on chromosome 5A, *ZCCT-B1* and *ZCCT-B2* on 4B and *ZCCT-D1* and *ZCCT-D2* on 4D, all on the distal long arm of the chromosomes (**Figure 1A**). The distance between these genes also differs between chromosome pairs, raising the possibility that *ZCCT* genes are under independent regulation and possibly even selection. It also highlighted that the individual genes of the *VRN2* loci could be non-redundant with respect to the vernalization response.

**Figure 1:**
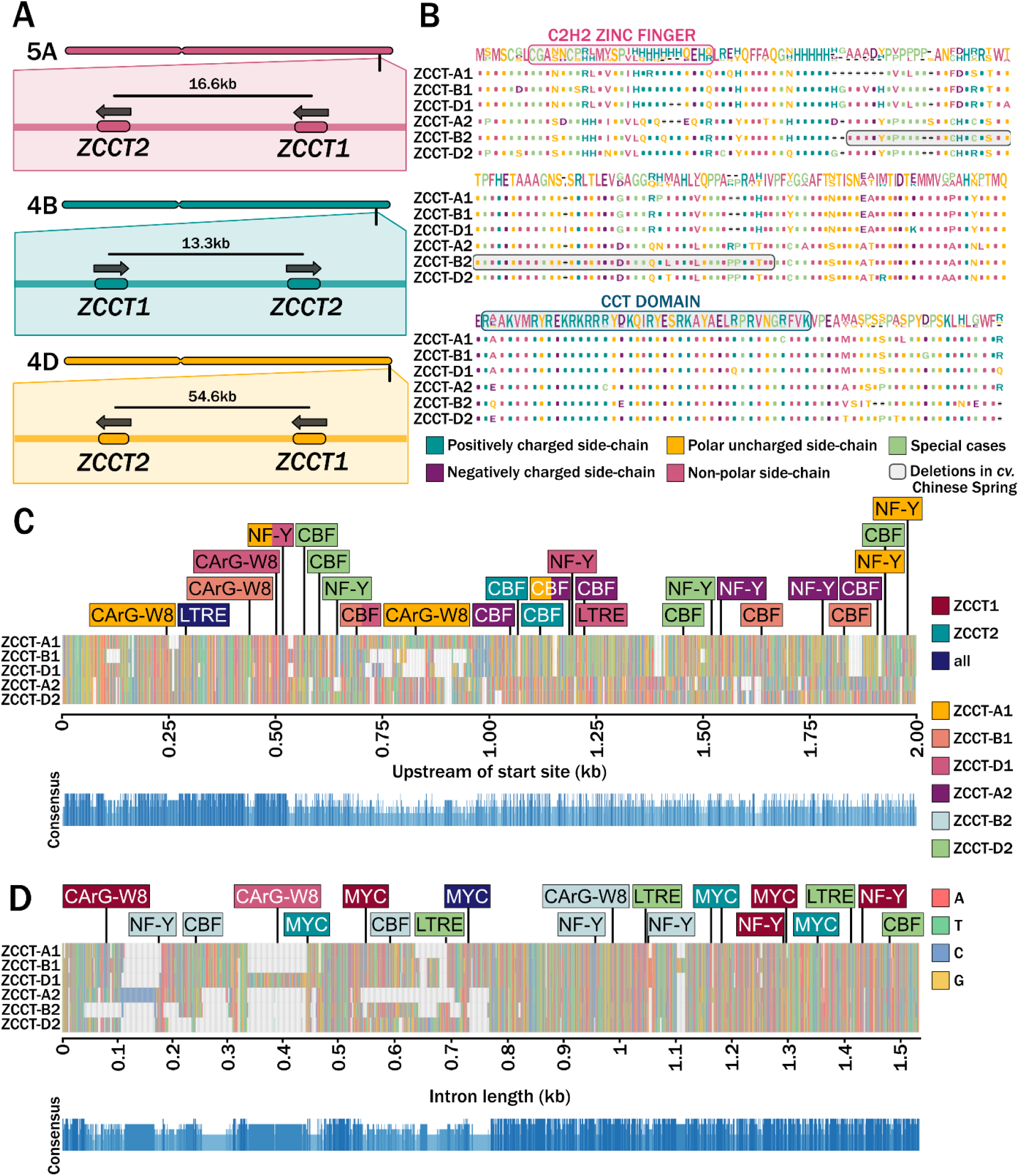
VRN2 location and sequence analysis. A) A schematic showing the direction and locations of each *ZCCT1* and *ZCCT2* gene on each chromosome, A, B and D as measured from the midpoint of each gene. B) The aligned protein sequences for each homoeologous copy of ZCCT1 and ZCCT2, highlighting the putative C2H2 zinc finger (pink) and CCT domain (blue). C) A consensus sequence indicates the level of conservation across the 5 ZCCT genes annotated in the v2.1 reference cultivar ‘Chinese Spring’ for the promoter region, 2kb upstream of the start site. Motifs were identified using ‘PlantPAN 4.0’ [23]. Nucleotides are indicated based on colour shown in the key and motifs of interest are indicated, with colours indicating whether the motif is conserved across all three homoeologous copies of ZCCT1 (maroon), ZCCT2 (teal) or across all ZCCT1 and ZCCT2 promoter regions (navy). Motifs which are specific to only one ZCCT promoter region are also indicated as indicated in the key. D) as for C, but for the intron.

The role of each *ZCCT* gene varies across different wheat species, with the diploid *T. monococcum* showing high expression of *ZCCT1* and low levels of expression of *ZCCT2*. In this wheat species, *zcct1* mutants exhibit a spring phenotype despite the presence of a functional *ZCCT2* gene, leading to the conclusion that *ZCCT1* is the primary functional copy [13]. In contrast, the tetraploid wheat species *T. turgidum* shows higher expression of *ZCCT2* than *ZCCT1* [12]. This species was used to generate a synthetic hexaploid wheat with a triple knock-out of *VRN2* [11], where the D-genome was contributed by *A. taushchii*. In this synthetic hexaploid, *ZCCT-B2* was identified as the key copy giving *VRN2* its function as a floral repressor [11]. This triple-null has a spring wheat phenotype, with significantly varying flowering times dependent on combinations of functional *VRN2* genome copies [11]. Understanding the role of each *VRN2* gene in natural hexaploid wheat will be of great interest, as this suggests the potential to modulate the vernalization requirement of wheat through allelic and copy number variation of specific *VRN2* genes.

Alongside monitoring ambient temperature to direct floral transition, photoperiod monitoring is integral to ensure floral transition is correctly timed. In both barley and rice, *VRN2* expression is regulated by photoperiod in conjunction with temperature [15, 16]. Photoperiod monitoring genes such as *PHOTOPERIOD1 (Ppd-1)* are part of a complex network of gene expression which regulates the central floral integrator, *FT1*, to ensure floral transition occurs when conditions are ideal [17]. Many of the genes with a role in monitoring photoperiod, including *Ppd-1* and the *CONSTANS (CO)* genes, also contain the highly conserved CCT domain [18-20]. In VRN2, mutations in the arginine amino acids at 16, 35 and 39 in the CCT domain significantly alter protein properties and can render the protein non-functional [12].

We asked how vernalization genes respond in facultative spring wheat under different environmental conditions and if plant development could be altered to improve winter traits without impacting the vernalization response. We identified that the expression of the *VRN2* loci is regulated by both temperature and photoperiod in bread wheat, and this expression alters under variable conditions. Therefore, specific genes within the *VRN2* loci can be targeted for improving aspects of crop performance. In particular, we identify that altered function of *ZCCT-D1* and -*D2* regulates early tiller outgrowth without impacting final tiller number and flowering time. This has the potential to improve water, light and nutrient efficiency as well as improve overall soil health and structure [7, 8, 21].

## Results

### ZCCT1 and ZCCT2, which form the VRN2 loci, contain different regulatory domains

The number of genes forming the *VRN2* loci in hexaploid bread wheat was believed to be six, *ZCCT-A1* and -*A2, -B* and *-D*. However, a blast analysis of the reference genome *cv*. Chinese Spring (v2.1) identified only five genes, missing *ZCCT-B2*. Specific gene search in the region of *ZCCT-B2* identified a truncated version of *ZCCT-B2* in the reference, containing a 62 amino acid deletion (**Figure 1B**). This highlights that the different *ZCCT1* and *-2* genes could be selected independently despite being physically close on the chromosome (**Figure 1A**). To start to understand this possibility we classified the regulatory domains within the intron and 2 kb upstream of the putative start site for each of the *ZCCT* genes (**Figure 1C** and **Figure 1D**). Here we could identify that the two genes *ZCCT-1* and *-2* contained unique regulatory domains but also that even within the *ZCCT-1* or *-2* genes independent selection had occurred. For example, in the 2 kb upstream of the putative start site a 0.25 kb region was deleted in *ZCCT-B1* and *-D1* compared with *ZCCT-A1*. There was also variation between genes for key motifs such as the CArG box, a binding site for MADS box transcription factors such as VRN1 [22, 23], which was present in all three homoeologous copies of *ZCCT1* in different locations but not *ZCCT2* (**Figure 1C**). The analysis of the single intron showed conservation was better in the latter half of the intron (approximately 0.8 kb – 1.5 kb) with large deletions present in *ZCCT-A2* and *ZCCT-B2* and insertions present in *ZCCT-D1* and *ZCCT-A2*, as well as several deletions in all three homoeologous copies of *ZCCT2* compared with *ZCCT1* (**Figure 1D**). This highlighted that the two *ZCCT* genes have diverged in regulatory domains and strongly suggested *ZCCT1* and *ZCCT2* are not functionally redundant and may be differentially regulated in response to environmental conditions.

### Core vernalization genes expression alters depending on temperature and photoperiod conditions in facultative wheat

The domain analysis suggested that the *VRN2* genes could be under different regulation and so we hypothesised that some of the *VRN2* genes may offer a route to coordinate environmental responses and development without always impacting the vernalization response. It has already been established that expression of the *VRN2* genes is responsive to changes in photoperiod [24], so to investigate the effect of temperature, we firstly characterized how vernalization genes behave in facultative spring wheat via RT-qPCR analysis on 3-week-old leaf tissue across a series of simulated environmental scenarios, all with the same day-neutral (DN; 12 h light:12 h dark) photoperiods.

We measured *VRN1* as this is the central regulator of vernalization and its expression is known to increase under lower ambient temperatures [4]. We observed a stable level of *VRN1* expression across the constant 10°C time course (**Figure 2A**). We then compared this with *VRN1* expression at constant 16°C as this temperature is associated with early season conditions in Northern Europe. Expression showed diurnal control with a peak before dusk (**Figure 2B**) and generally higher expression than under constant 10°C. Given that *VRN1* is primarily regulated by cold exposure and not traditionally considered under circadian control, it was unexpected to observe rhythmic expression under warmer conditions.

**Figure 2:**
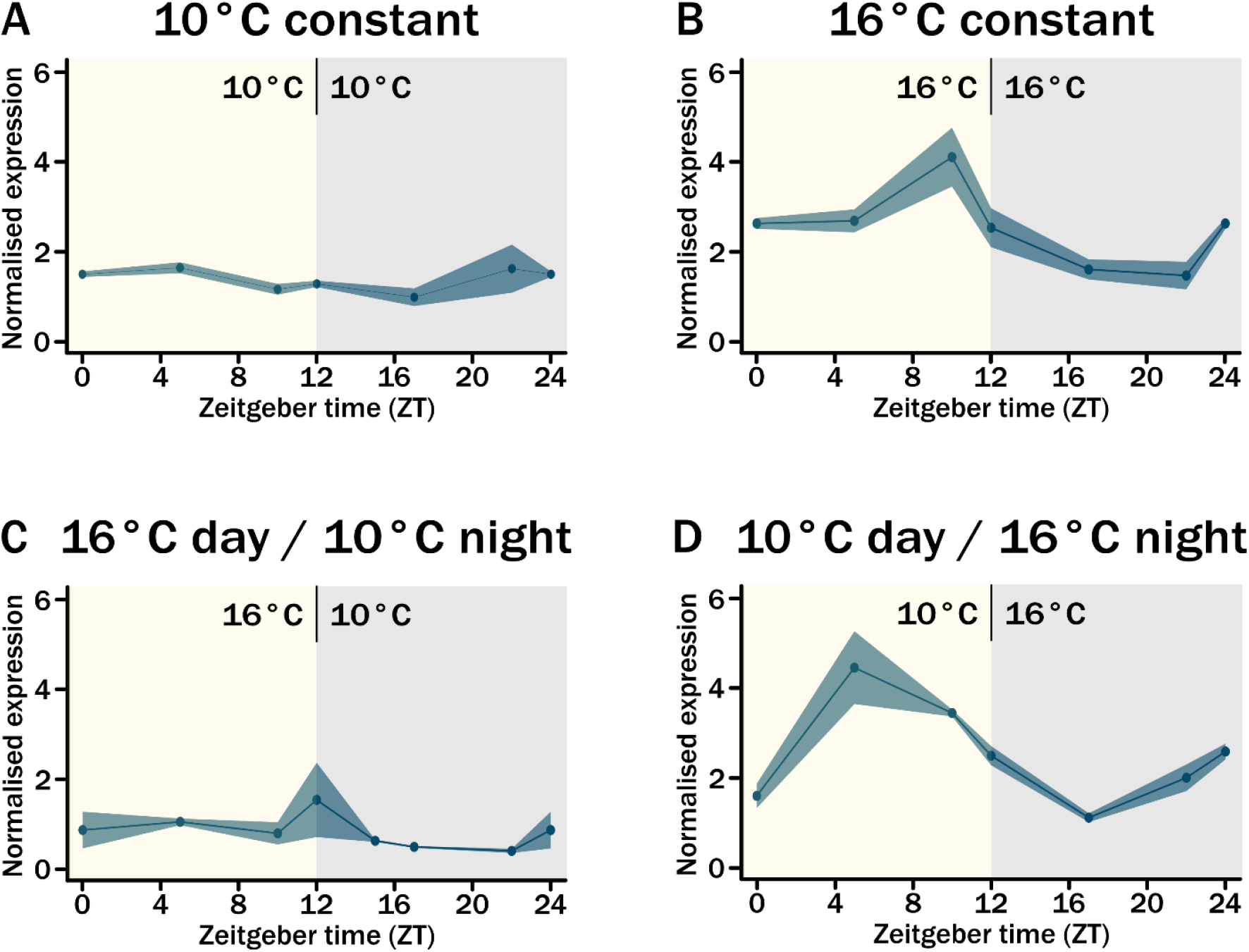
Expression of *VRN1* under day-neutral conditions with variable temperatures. A ribbon plot showing expression of *VRN1* for all three genome copies (A,B,D) across a 24-hour time course. Leaf tissue was sampled after 3 weeks of growth of spring ‘Cadenza’ plants in 12-hour day-neutral conditions under different temperatures. Expression was normalised against *TraesCS5A02G015600* and n=3 for each time point. Standard error mean (SEM) is indicated by the shaded region around each point. The light period is indicated by yellow and dark period by grey background shading. ZT0 is the same sample as ZT24 except for D. A) Expression of *VRN1* under constant 16°C conditions. B) as for A but for constant 10°C conditions. C) as for A but for 16°C light / 10°C dark conditions. D) as for A but for 10°C light / 16°C dark conditions.

The expression patterns of constant 10°C and 16°C suggested that the constant 10°C conditions were actually impacting the day expression pattern of *VRN1*, to investigate this we conducted the 24 h time course under 16°C by day and 10°C at night DN conditions. Here, we again observed a low, constant level of *VRN1* expression with a slight hint of a pre-dusk increase in expression (**Figure 2C**). This indicated that in the facultative spring wheat it was actually the night temperature which was driving a diurnal expression of *VRN1*. To challenge this, we conducted a final 24 h time course during which we altered the day and night temperatures to generate an unrealistic 10°C by day and 16°C at night DN time course. Here the diurnal rhythmicity is returned with a peak during the day (**Figure 2D**). This highlights that in the facultative spring wheat, Cadenza, the expression of the main vernalization gene is regulated by the interaction of photoperiod and temperature, where previously it has only been considered temperature responsive. Furthermore, it highlights a role for warming night temperatures in regulating its expression during the subsequent day.

As a main regulation target of VRN1 is the repression of *VRN2* [25], we anticipated that the temperature dependent expression patterns observed for *VRN1* would also be reflected through the regulation of *VRN2*. Measuring the two *VRN2* genes, *ZCCT-1* and *-2*, as homoeologous groups across the previously described 24 h time courses we observed quite distinct patterns of regulation. Under constant 10°C DN both *ZCCT-1* and *-2* expression was highly reminiscent of the low, constant expression observed for *VRN1* (**Figure 3A**). However, unlike *VRN1* under constant 16°C DN both *ZCCT-1* and *-2* still had a low level of expression (**Figure 3B**). Only when considered on a higher resolution scale, due to the low actual expression levels, could *ZCCT-2* be identified to show a similar diurnal pattern to that observed for *VRN1* (c.f. **Figure 2A** and **Figure 3A**) whilst *ZCCT-1* lacked this rhythmicity. This supported previous reports that the *ZCCT* genes are only expressed under longer photoperiods [15, 24]. However, when we measured *ZCCT-1* and *-2* expression under changing light/dark temperature conditions both were expressed and showed a diurnal pattern in expression, which was synchronous between the genes (**Figure 3C**). Notably, depending on the temperature pattern the peak in expression shifted such that under warm days the peak was across and just after dusk whilst with warm nights it was during the light period. Under these conditions the expression of *VRN2* is asynchronous with *VRN1*. Furthermore, through comparing the four 24 h time courses it identified that temperature is an important regulator in the expression of these genes and that the interaction between the genes may be more complex as the patterns of their expression are altering with variable conditions. We therefore aimed to understand how the vernalization genes expression was responding across a growing season.

**Figure 3:**
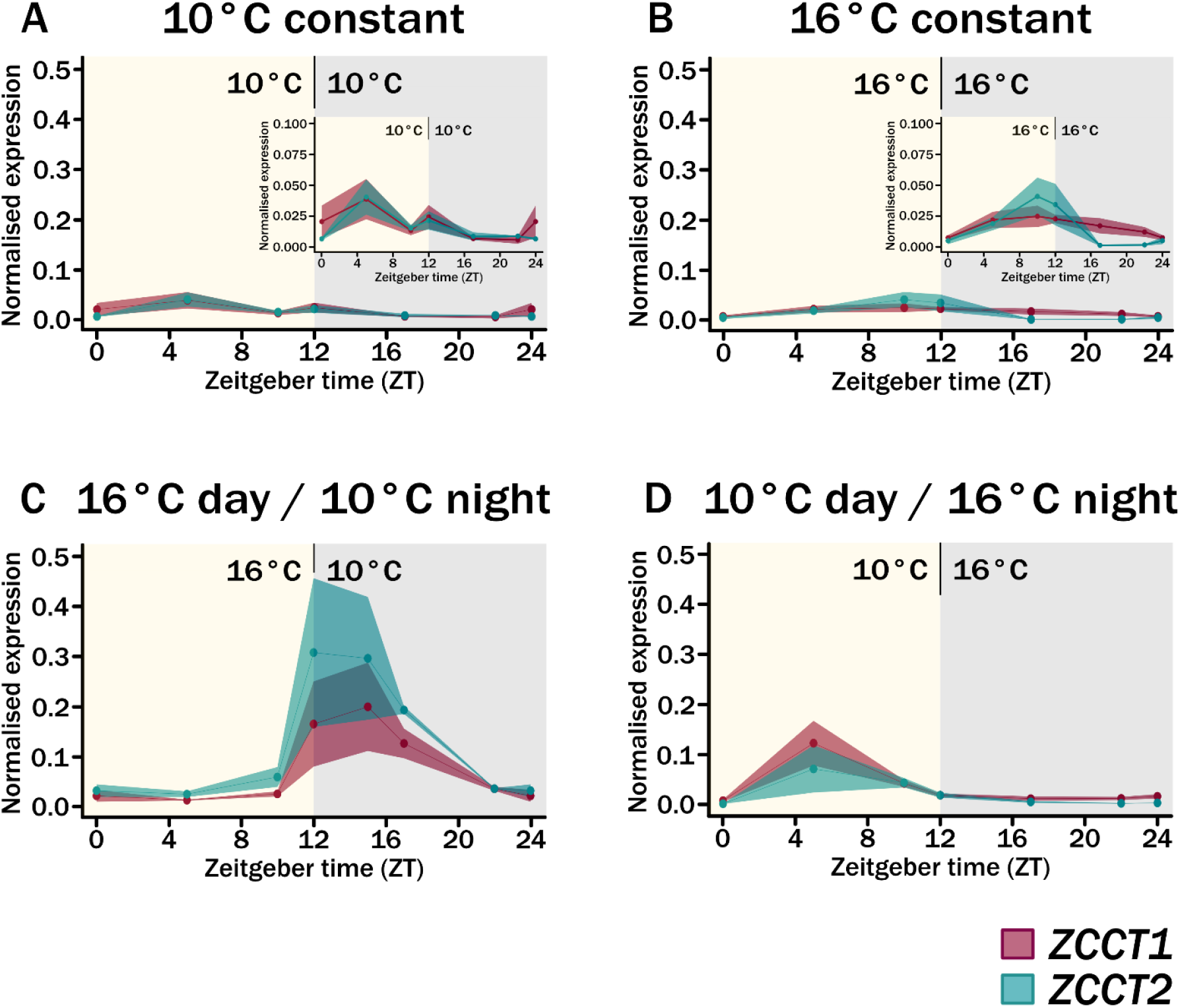
Expression of *ZCCT1* and *ZCCT2* under day-neutral conditions with variable temperatures. A ribbon plot showing expression of *ZCCT1* for all three genome copies (A,B,D) and *ZCCT2* (A,B,D) in pink and teal respectively across a 24 hour time course. Leaf tissue was sampled after 3 weeks of growth of spring ‘Cadenza’ plants in 12-hour day-neutral conditions under different temperatures. Expression was normalised against *TraesCS5A02G015600* and n=3 for each time point. Standard error mean (SEM) is indicated by the shaded region around each point. The light period is indicated by yellow and dark period by grey background shading. ZT0 is the same sample as ZT24 except for D. A) Expression of *ZCCT1* and *ZCCT2* under constant 16°C conditions. B) as for A but for constant 10°C conditions. C) as for A but for 16°C light / 10°C dark conditions. D) as for A but for 10°C light / 16°C dark conditions.

### ZCCT1 expression increases with photoperiod post-vernalization

The role of varying day and night temperatures in the regulation of the vernalization genes expression suggested that both environmental conditions play an important role in the regulation of these genes. It also indicated that even in a facultative spring wheat the role of cold was important for the synchronisation of gene expression. We therefore wanted to test this across a more realistic time course. To do this we generated an experimental profile where we took the approximate annual environmental conditions for London, UK. We calculated the average day-length, temperature during the light-period and temperature during the dark-period for each calendar month to generate a time course over 12 weeks with each week containing the average environmental condition for one calendar month. We added in one more week of the mid-winter condition to ensure that the plants experienced a true vernalization period in case our experimental time course was not sufficient (**Figure 4A**). Cadenza seeds were stratified and then grown in soil from day 0 of week 1. Under these conditions Cadenza flowered following 125 days ±1.73 SD, which confirmed that the conditions were sufficient to support floral development.

**Figure 4:**
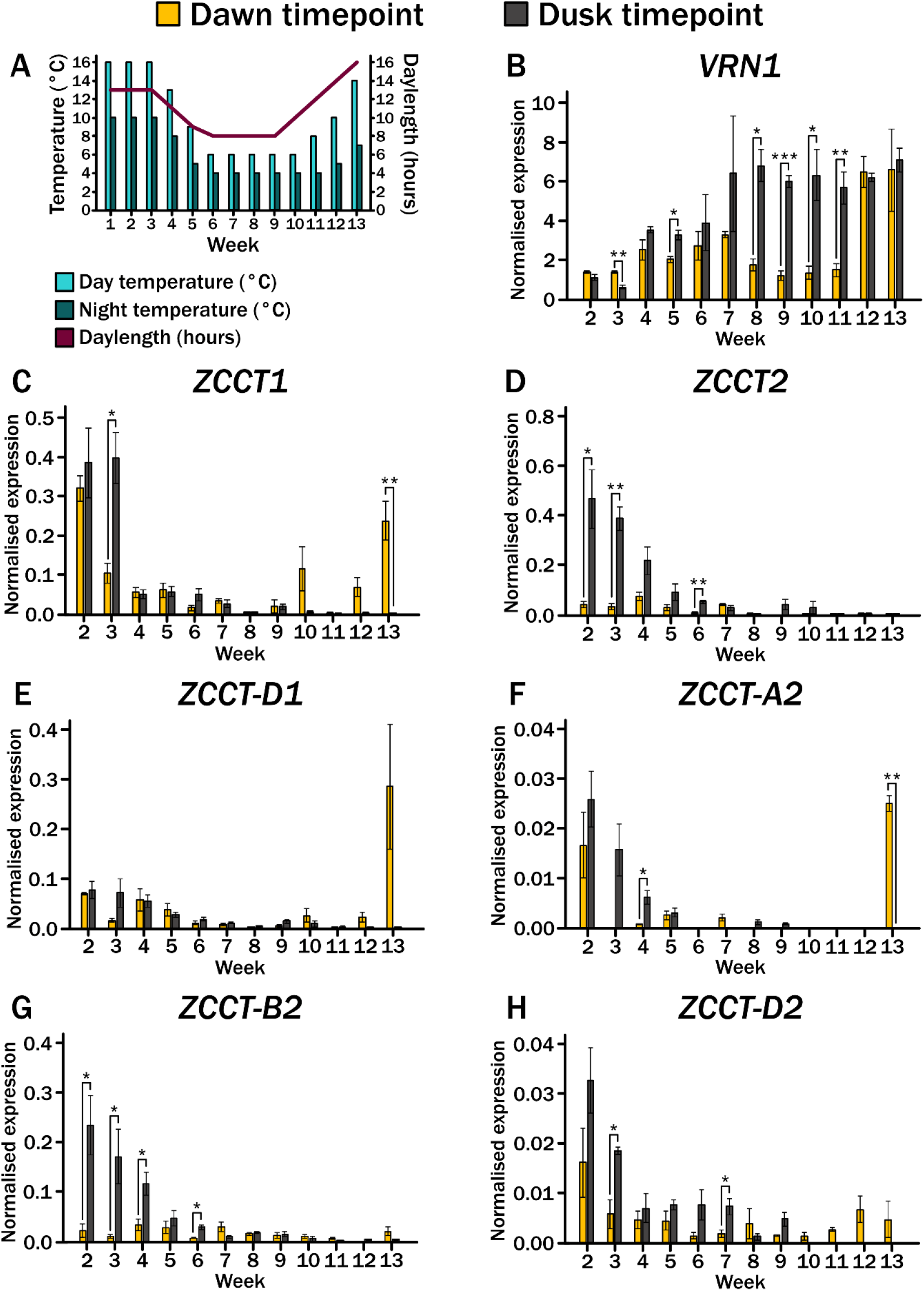
Gene expression across the long-term seasonal time course at dawn and dusk for the facultative spring wheat ‘Cadenza’. A) A graph outlining the temperature conditions for each week during the day (yellow) and night (grey), as well as the daylength indicated by number of hours of light (red). B) Bar graphs showing gene expression across the long-term seasonal time course for facultative spring wheat ‘‘Cadenza’. Leaf tissue was sampled 1 hour after the relative ‘dawn’ and ‘dusk’ time points and n=3 for each time point. Expression of *ZCCT1* was normalised against *TraesCS5A02G015600* and standard error mean (SEM) is indicated by the error bar. \**P<0*.*05*, \*\**P*<0.01, \*\*\**P*<0.001 determined by paired Student’s *t*-test. C) As for B, but for expression of *ZCCT2*. D) As for B, but for expression of *VRN1*. E) As for B, but for expression of *ZCCT-D1*. F) As for B, but for expression of *ZCCT-A2*. G) As for B, but for expression of *ZCCT-B2*. H) As for B, but for expression of *ZCCT-D2*.

To assess the gene expression, leaf tissue from the newly emerging leaf was sampled each week one hour after dawn and dusk. Confirming the observations from the 24 h time course (**Figures 2 and 3**) gene expression of *VRN1* and *2* differed at these two time points. *VRN1* showed a steady increase across the time course, as previously reported [26, 27], consistent with its role in floral activation (**Figure 4B**). However, and of possible equal biological interest is the substantial, over 3-fold increase between week eleven to twelve observed at the dawn time point. This coincides with an increasing day temperature from 8°C to 10°C, night temperature from 4°C to 5°C and photoperiod 12 to 14 as well as representing the end of the vernalization period.

The *ZCCT* genes also showed differences in expression between the dawn and dusk time points (**Figure 4C-H**). For both *ZCCT-1* and *-2* the dusk time point showed a steady decrease in expression as photoperiod and temperature decreased from weeks two to five. Expression then remained largely repressed across the time course (**Figure 4C-H**), this is consistent with its repression as a result of vernalization [13]. However, like *VRN1* the same pattern was not observed at the dawn time point. Here the genes not only showed a different regulation to dawn but also between the two *ZCCT* genes. For *ZCCT-1* expression also decreased at dawn with the decreasing temperature and photoperiod but it was not robustly repressed when temperature and photoperiod started to increase again, at week ten onwards (**Figure 4C**). This is in contrast to *ZCCT-2* which was also sequentially repressed across the time course at the dusk time point as temperature and photoperiod decreased but was hardly expressed at the dawn time point (**Figure 4D**).

As *ZCCT-1* and *-2* showed different expression responses across the seasonal time course we next asked if these differences extended to the level of homeologous genes. To assess this, we designed gene specific RT-qPCR primers. Due to the high level of homology between the genes we were only able to design primers which were specific for each of the *ZCCT-2* genes and *ZCCT-D1* (details provided in **Supplementary Table 1**). Between the *ZCCT-2* genes there were distinct expression patterns, with *ZCCT-A2* and *-D2* (**Figures 4F** and **4H**) showing an increase in expression following the winter-simulated period whilst *ZCCT-B2* did not (**Figure 4G**). This trend was not observed using the generic *ZCCT-2* primers, possibly because the expression level of *ZCCT-A2* and *-D2* was very low (10-fold lower). *ZCCT-D1* expression followed that of the generic primers, although its expression was lower than the generic primers at the start of the time course (**Figure 4E**). For all of the genes there were differences in expression between the dawn and dusk time points.

### Early tiller development is accelerated in VRN-D2 TILLING lines

The expression of the *ZCCT* genes and sub-genome copies suggested that in facultative spring wheat they can function beyond the vernalization response. Therefore, we aimed to identify if any of the *VRN2* genes controlled important developmental traits without significantly altering the facultative spring vernalization response. To do this we utilised the Cadenza TILLING collection [28] and were able to identify TILLING mutants which contained a SNP within the coding sequence of two of the *ZCCT* genes. The line *Cadenza0810* contained a predicted stop Q144* just before the CCT domain in *ZCCT-D2*, hereafter *zcct-d2_m1*, whilst *Cadenza1436* contained a T130I, also preceding the CCT domain in *ZCCT-1D*, hereafter *zcct-d1_m1* (**Figure 5A**).

**Figure 5:**
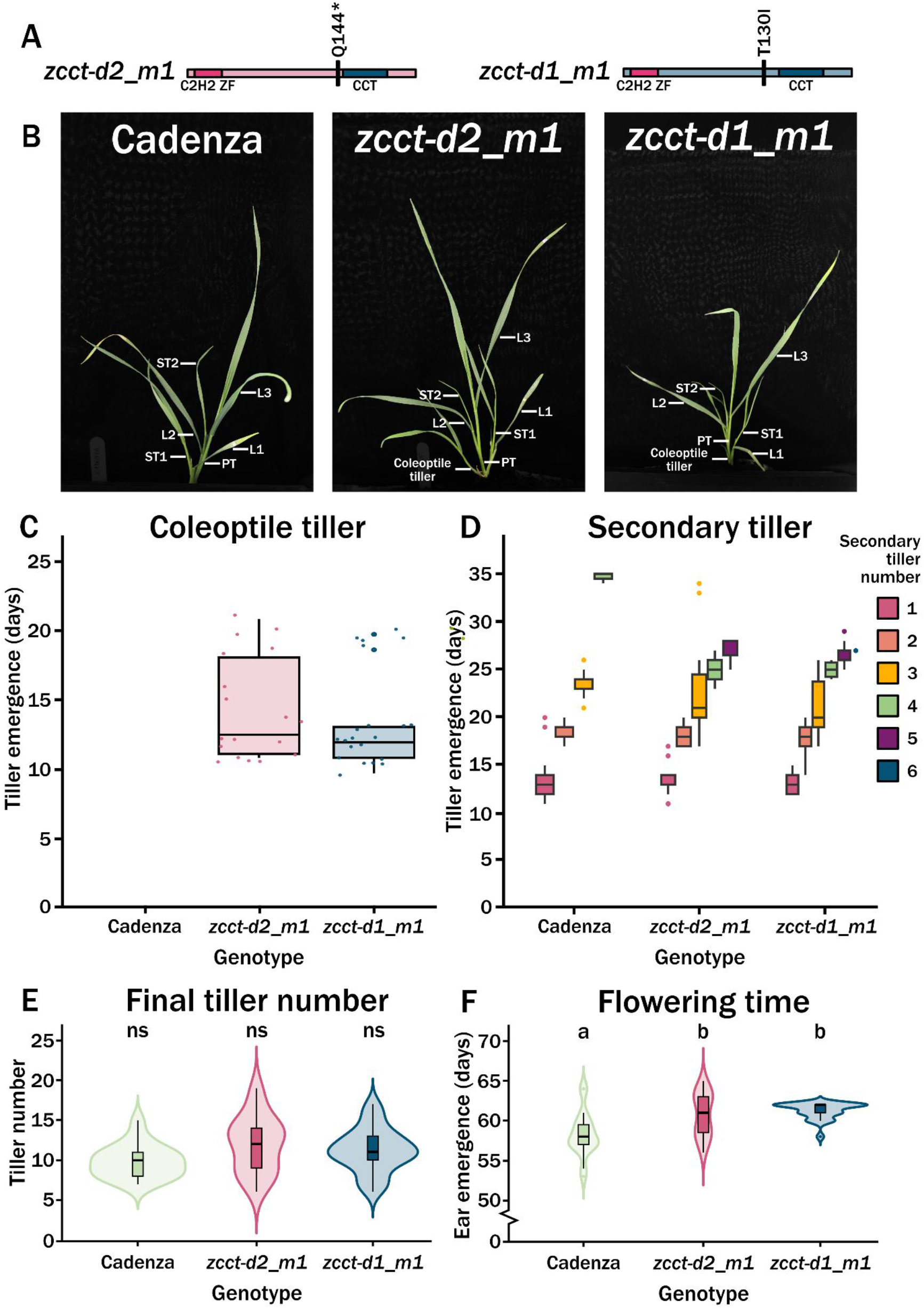
Tiller growth in *VRN2* mutants. A) An infographic outlining the locations of the mutations in the TILLING lines used in this study, *zcct-d2_m1* with a premature termination codon at Q144*, and *zcct-d1_m1* with a T130I missense mutation. All germplasm used was at the BC_2_F_2_ stage, homozygous for the mutations shown. The relative positions of the C2H2 zinc finger (pink) and CCT (blue) domains are also as illustrated. B) A labelled image of example wheat plants for the ‘Cadenza’ control, *zcct-d2_m1* and *zcct-d1_m1* with the primary tiller (PT) and coleoptile tillers indicated, along with the emerging leaves (L1, L2, L3) and secondary tillers (ST1, ST2) numbered in ascending order. C) Box plots showing the mean average time in days until emergence of a new coleoptile tiller in the ‘Cadenza’ control (n=19) compared with the two independent mutant lines; *zcct-d2_m1* (n=21) and *zcct-d1_m1* (n=24). D) as for C, but for emergence of secondary tillers, with colours indicating the secondary tiller number (1=pink, 2=orange, 3=yellow, 4=green, 5=purple, 6=blue). E) Violin plots showing the final tiller number, \**P<0*.*05*, \*\**P*<0.01, \*\*\**P*<0.001 determined by a Kruskal-Wallis test followed by a pairwise Wilcoxon rank sum test with Bonferroni correction. F.) as for E, but for flowering time defined as half-ear emergence (GS55).

To reduce the possible impact of background mutations each line was backcrossed twice and homozygous mutations were selected to generate homozygous BC_2_F_2_ lines. These lines were grown under 16°C LD conditions to replicate conditions at the start of autumn and the plants were carefully phenotyped. Noticeably, a coleoptile tiller developed in the two TILLING mutants but not Cadenza (**Figure 5C**). Then across the first 30 days of growth both *zcct-d2_m1* and *zcct-d1_m1* developed an additional two to three tillers than Cadenza (**Figure 5D**). This increase in early tiller development was not maintained and final fertile tiller number was similar between all lines (**Figure 5E**). The increase in early tiller development was however reflected by a slight, but significant (p=0.01), delay in flowering time of 2.5 days ± 2.74 SD for *zcct-d2_m1* and 2.9 days ± 1.03 SD for *zcct-d1_m1* as measured by half spike emergence (Waddingtons GS55; **Figure 5F**). This slight delay did not affect the spikelet number (**Supplementary Figure 1**).

### ZCCT genes co-regulate their expression

As we observed an early stage developmental phenotype we next asked if this altered the expression of the vernalization genes. To test this, we employed our condensed season time course, as described for **Figure 4A**. We sampled emerging leaf tissue at dusk and measured the expression of vernalization genes in Cadenza, *zcct-d2_m1* and *zcct-d1_m1*. Here we observed that in plants carrying mutant alleles of *ZCCT-D1* or *-D2 VRN1* expression was higher, and this was particularly apparent at the start of the time course (weeks one and two) (**Figure 6A**). This increase in *VRN1* was also reflected by a generally lower expression level of *ZCCT-1* and *-2* as measured using the generic primers for these genes (**Figure 6B** and **6C**). This supports the molecular inter-regulation of these genes, as previously described [13, 25]. Interestingly, this general trend was not maintained when we measured the expression of individual *ZCCT-1* and *-2* genes. Here we observed that the function of the *ZCCT*s impacted the expression of other *ZCCT* genes. The expression of *ZCCT-A2* was increased in both lines which carried either less functional *ZCCT-D1* or *-D2*, suggesting that usually these genes repress the expression of *ZCCT-A2*. Similarly at the start of the time course *ZCCT-D2* expression was much lower in the *zcct-d1_m1* suggesting that *-D2* is activated by *-D1*. The reverse was observed in the *zcct-d2_m1* which had higher expression of *-D1*, highlighting a possible mutual regulation mechanism (**Figures 6D** and **6G**). Significant changes in gene expression were most apparent at the start of the time course, supporting a role for these genes in early developmental regulation. Additionally, we measured flowering times and observed a flowering delay in the *zcct-d2_m1* line, but not for *zcct-d1_m1* (**Figure 6H**).

**Figure 6:**
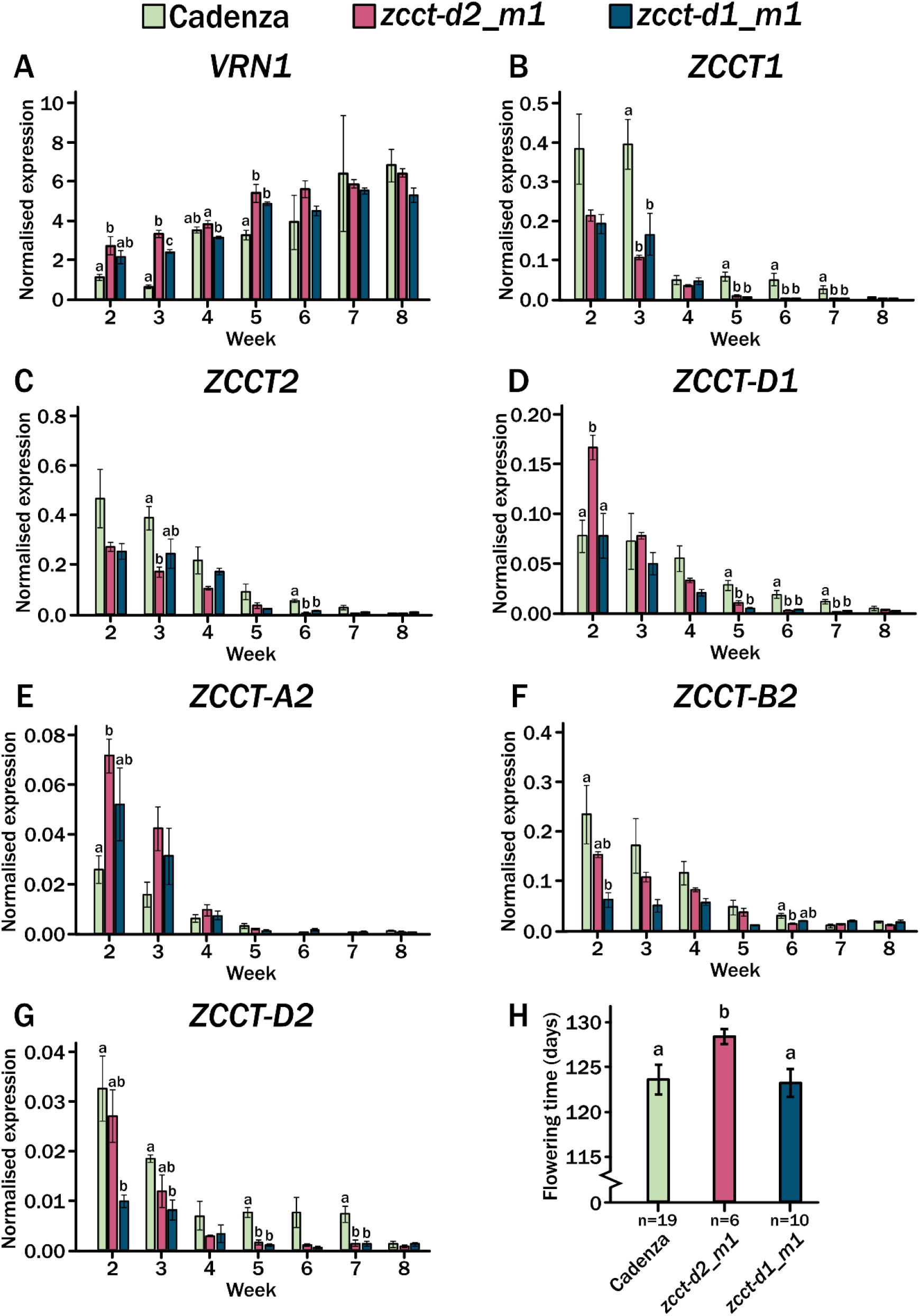
Gene expression across the long-term seasonal time course for the facultative spring wheat ‘Cadenza’ and mutant lines *zcct-d2_m1* and *zcct-d1_m1*. Bar graphs showing gene expression across the long-term seasonal time course up to week 8 for facultative spring wheat ‘Cadenza’ (green) and mutant lines *zcct-d2_m1* (pink) and *zcct-d1_m1* (blue). Leaf tissue was sampled 1 hour after the relative ‘dusk’ time point and n=3 for each time point. Expression was normalised against TraesCS5A02G015600 and standard error mean (SEM) is indicated by the error bar. \**P<0*.*05*, \*\**P*<0.01, \*\*\**P*<0.001 determined by a one-way ANOVA followed by a Tukey’s Honestly Significant Difference post-hoc test. A) Expression of *ZCCT1*. B) Expression of *ZCCT2*. C) Expression of *VRN1*. D) Expression of *ZCCT-D1*. E) Expression of *ZCCT-A2*. F) Expression of *ZCCT-B2*. G) Expression of *ZCCT-D2*. H) The flowering time for each genotype (Cadenza in green, *zcct-d2_m1* in pink, *zcct-d1_m1* in blue) following growth in the outlined conditions. Error bars indicate standard deviation; n=19 for Cadenza, n=6 for *zcct-d2_m1* and n=10 for *zcct-d1_m1*.

## Discussion

### Vernalization genes are responding to multiple environmental factors

Vernalization is the requirement for cold to enable the vegetative to floral transition and this definition reflects the absolute need for lower ambient temperatures during this process [4]. Additionally, as a seasonal response photoperiod plays an important role, in particular regarding the repression of *VRN2* expression by short-days [15, 24]. With the increasingly variable and unpredictable conditions of winter we aimed to further understand how temperature and photoperiod interacted to regulate the expression of the major vernalization genes in hexaploid bread wheat. We were particularly interested in understanding this function in facultative spring wheat as this growth habit offers a mechanism of increased flexibility to regulate plant development under variable winters. We measured *VRN1* and *VRN2* expression under different temperature patterns with day-neutral photoperiods to further our understanding of how temperature influences the expression of these genes. Interestingly, we observed that both *VRN1* and *VRN2* had altered expression patterns depending on the temperature (**Figures 2 and 3**). Of particular interest is the altered waveform in expression which occurred in response to different temperature conditions. This is highly reminiscent of what is observed in barley (*Hordeum vulgare*) [29]. It was also notable that the circadian regulation is more robust for *VRN1* under warmer temperatures (**Figure 2B**). Furthermore, *VRN2* indicated a possible response to temperature entrainment, such that its expression altered significantly when temperature cycles were introduced and that the phase of the expression altered depending on the cycle temperatures. This is an interesting response which could hint at possible reasons for altered vernalization responses observed depending on quite small temperature changes, or when major day/night temperature differentials are observed. For example, winters which have greater day/night differences in temperature may take longer for plants to vernalize due to the activation of *VRN2* expression at night. This would be compounded by our observation that *VRN1* expression is greater during the day. Therefore, under certain environmental conditions *VRN1* and *VRN2* expression is asynchronous, suggesting other genetic factors are involved in the regulation of these genes.

### Vernalization genes show different responses across development

To better understand how these genes respond under natural conditions, we conducted a seasonal time course. The possibility of multiple factors regulating the vernalization gene expression patterns appears more apparent in the seasonal time course (**Figure 4**). Here, *VRN1* expression differs between dawn and dusk time points. At dusk, its expression follows the reported trajectory of increasing during and following vernalization. In contrast, at dawn, expression shows a gradual increase, followed by decrease and then sudden increase post-vernalization as spring conditions are reached. This strongly suggests that the gene is being repressed at dawn until post-vernalization conditions. Similarly, the *VRN2* genes displayed differential expression patterns between dawn and dusk conditions. Notably for *ZCCT-D1* and *-A2* expression was not robustly repressed post-vernalization, suggesting that these genes may have additional roles in regulating wheat development after the floral transition has occurred. These expression patterns strongly imply that the *VRN2* genes have additional roles beyond the timing of the apex transition and that future work dissecting their function in later stage development would be very interesting. Notably, *VRN1*, in conjunction with *FUL2* and *3* has an important role in apex patterning and spikelet development [30] and so additional roles for the *VRN2* genes could be anticipated.

### Early tiller growth is regulated by ZCCT-D1 and -D2

We aimed to characterise the vernalization genes in a facultative spring wheat and understand if through considering the *VRN2* genes separately, we could identify routes to alter winter-type traits without impacting the floral regulation. To test this hypothesis, we developed two mutant alleles in *ZCCT* genes and phenotyped them under 16C LD. We observed that the *ZCCT-D1* and *-D2* genes were involved in the regulation of lateral branch, or tiller, out-growth early in plant development (**Figure 5C** and **5D**). This was further supported when we measured the expression of these genes across our condensed growing season time course. Here we observed differences in gene expression in the first few weeks of the time course which returned to a similar point by week five (**Figure 6**). Furthermore, the final plant phenotypes were extremely mild, with the final tiller number the same between all lines, along with spikelet number (**Figure 5E** and **Supplementary Figure 1**). The absence of major developmental phenotypes at the later growth stages suggests that changes in ZCCT gene expression did not significantly impact overall plant growth. However, we did observe a small flowering time change with the mutant alleles flowering approximately two days later under constant conditions and 5 days later under variable conditions for *zcct-d2_m1* which is not the expected result for a floral repressor gene (**Figures 5F** and **6H**). This suggests that the function of the individual *ZCCT* genes differs and that their role in plant development may be quite complex when considered outside of the scope of vernalization.

### ZCCT genes show co-regulation

Unexpectedly, when we measured the individual gene expression in the mutant backgrounds we observed that some of the *ZCCT* genes appeared to regulate other *ZCCT* gene expression. For example, between *ZCCT-D1* and *-D2* a regulation loop could be inferred where a less-functional *-D1* caused a reduction in expression of *ZCCT-D2*, suggesting that normal functional *-D1* promotes the expression of *-D2* (**Figure 6G**). Whereas, the expected less functional *-D2* led to higher *ZCCT-D1* expression, suggesting its normal role is to repress the expression of *-D1*, however, only at the very start of the time course (**Figure 6D**). Furthermore, the genetic and expression data also indicate that both *ZCCT-D1* and *ZCCT-D2* repress the expression of *ZCCT-A2*, and that this occurred throughout the vernalization period of the time course (**Figure 6E**). Therefore, the gene specific analysis has identified that the different *ZCCT* genes are not only not all expressed under the same regulation (**Figure 6**) but also are able to regulate their own expression. This is a common regulatory mechanism within adaptive response genes [31] but has not been previously reported for cereal vernalization.

Here we show that the core vernalization genes in facultative spring hexaploid wheat cv. Cadenza are regulated differently depending on the temperature conditions experienced and that this regulation causes an asynchronous expression pattern between the main repressor and activator in the vernalization pathway. Furthermore, when considering each of the individual *ZCCT* genes we identified inter-regulation in expression as well as independent regulation. Through the development of germplasm which differed between *ZCCT* genes we were able to identify variation in early development of tiller growth, which may have applications in the use of facultative spring wheat. It is a beneficial phenotype due to the increased coverage during early development which can improve water, light and nutrient efficiency as well as improve competition with biotic stressors [7, 9, 21]. To understand the potential use of these mutants, they will need to be transferred to current breeding material and tested for early development ground cover in field trials.

Exploring the roles of specific *ZCCT* genes has shown that these genes have further adaptive potential not only in the context of vernalization but potentially in other developmental roles. This means these genes may also be involved in regulating development in wheats which do not have a winter habit, offering the potential to regulate plant development without impacting vernalization per se.

## Materials and Methods

### Plant materials and growth conditions

All plants were germinated at room temperature (21°C) on Whatman filter paper with 5 ml ddH2O in a 9 cm petri dish and seedlings transferred to 24-well seed trays (each cell 50 mm x 48 mm x 52 mm) containing John Innes Cereal Mix [32]. The cultivars used were either from laboratory stocks (Cv. Cadenza) or obtained from the Germplasm Resource Unit (GRU) for all TILLING mutants [28]. Plants used for generation of *zcct-d2_m1* and *zcct-d1_m1* mutant lines were then grown under glasshouse conditions (16 hours light, 21°C/8 hours dark, 15°C).

The facultative spring wheat cultivar ‘Cadenza’ was used for the 24 h time course expression analysis. These plants were grown in Sanyo Plant Growth cabinets under the following temperature conditions with a constant 12-hour photoperiod; 16°C during the light period/10°C during dark, 10°C during light/16°C during dark, 10°C constant and 16°C constant. Plants were grown in these conditions for 3 weeks and then leaf tip tissue was sampled at 5-hour intervals across a 24-hour period, with tissue from 2 individual plants pooled for each of 3 biological replicates.

Growth conditions for the long-term seasonal gene expression experiment were conducted in a Conviron gen 2000 growth chamber with conditions listed in ***Supplementary Table S2***. The facultative spring wheat cultivar ‘Cadenza’ and mutant lines *zcct-d2_m1* and *zcct-d1_m1* were grown in these conditions. Leaf tissue samples were taken each week an hour after the relative ‘dawn’ and ‘dusk’ for each day depending on when light changes occurred.

Tiller emergence was recorded every day for the first 35 days of growth. Flowering time was defined as half-ear emergence from the flag leaf, GS55 on the Waddingtons scale. Spikelet number was counted for the spike from the primary tiller for each plant.

### Generation of zcct-d2_m1 and zcct-d1_m1 mutant lines

The two mutant lines were in the hexaploid ‘Cadenza’ background, Cadenza0810 and Cadenza1436 (with SNPs in *ZCCT-D2* and *ZCCT-D1* respectively) (Ks Tilling ref) were twice backcrossed. Plants were genotyped, using primers detailed in **Supplementary Table S1**, at the SNP of interest via KASP assay as described in [33] to identify homozygous plants. Genomic DNA was extracted using the chloroform:isoamyl alcohol method adapted from [34].

PCR amplification was conducted to confirm homozygosity at the SNP of interest using Q5 DNA Polymerase (NEB) with reagent quantities and conditions following the manufacturer’s protocol. Primers are listed in **Supplementary Table S1**. The PCR reaction was separated using gel electrophoresis and the fragment was extracted using the Monarch® DNA Gel Extraction Kit (NEB) according to the manufacturer’s protocol. Samples were then sequenced by Eurofins Genomics and analyzed to confirm the SNP.

### Analysis of gene expression

To measure the expression plants were sampled as outlined in *Plant materials and growth conditions*. All plant samples were flash-frozen in liquid nitrogen and stored at −70°C. The tissue was lysed using the TissueLyserLT (Qiagen) using 3mm steel ball bearings, and RNA was then extracted for all leaf tissue samples using the Spectrum™ Plant Total RNA Kit (Sigma-Aldrich) or Monarch® Total RNA Miniprep kit (NEB) according to the relevant manufacturer’s protocol. Synthesis of cDNA was conducted according to either [33] or using the reverse transcriptase UltraScript 2.0 (PCR Biosystems) according to the manufacturer’s protocol. The cDNA was then diluted (1:10) and quantitative real-time reverse transcriptase PCR (RT-qPCR) was conducted using GoTaq® qPCR Master Mix (Promega) according to the manufacturer. The primers used are listed in **Supplementary Table S1**, and the thermal cycling conditions performed using the CFX96™ Thermal Cycler (Bio-Rad) and protocol described in [33]. Expression was calculated relative to the housekeeping gene *TraesCS5A02G015600* [35], using the formula 2ΔCT where ΔCT= [expression of Gene of Interest] – [expression of *TraesCS5A02G015600*].

## Supplemental Materials

**Figure S1**. Final spikelet number for Cadenza, *zcct-d2_m1* and *zcct-d1_m1* in 16°C LD conditions.

**Figure S2**. Expression pattern of the individual genome copies of *ZCCT1* and *ZCCT2* under varying temperature conditions in a day-neutral photoperiod.

**Table S1**. List of primers

**Table S2**. Weekly growth conditions for long-term gene expression experiment

**Table S3**. List of germplasm

## Author contributions

D. H., and L.D. conceived, designed and conducted the experiments and analysis. H. T., I. L. and A. G. designed and conducted the experiments. D. H. and L.D. wrote the initial manuscript and all authors have confirmed its accuracy.

## Acknowledgements

We thank Dr Rachel Rusholme-Pilcher at Earlham Institute, UK for the assistance with the Cadenza *VRN2* gene chromosomal locations. We thank Dr Beth Soanes for her assistance in sampling the long-term seasonal time course. This research was supported through funding to L.E.D. via a UKRI FLF MR/S031677/1, Rank Prize Funds New Lecturer Award and start-up funds from the University of Leeds. A Molecules to Landscapes grant from BBSRC supported H.T. The authors have no conflicting interests.

