## Supplementary files for "*VERNALIZATION2 (VRN2)* alters early tiller development in a facultative spring hexaploid bread wheat (*Triticum aestivum*)"

**Supplemental Materials**

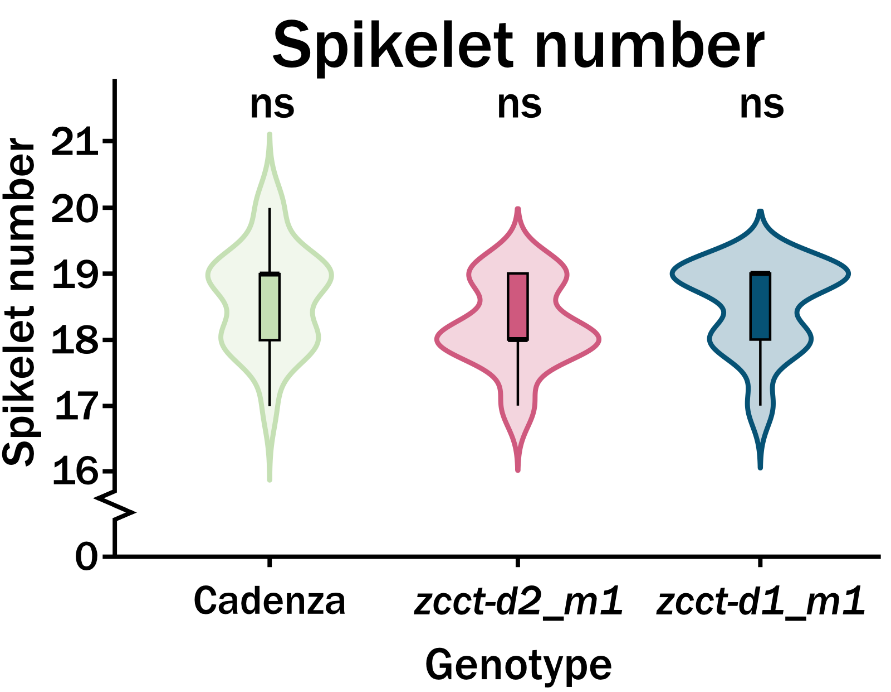

**Figure S1.** Final spikelet number for Cadenza, *zcct-d2_m1* and *zcct-d1_m1* in 16°C LD conditions.

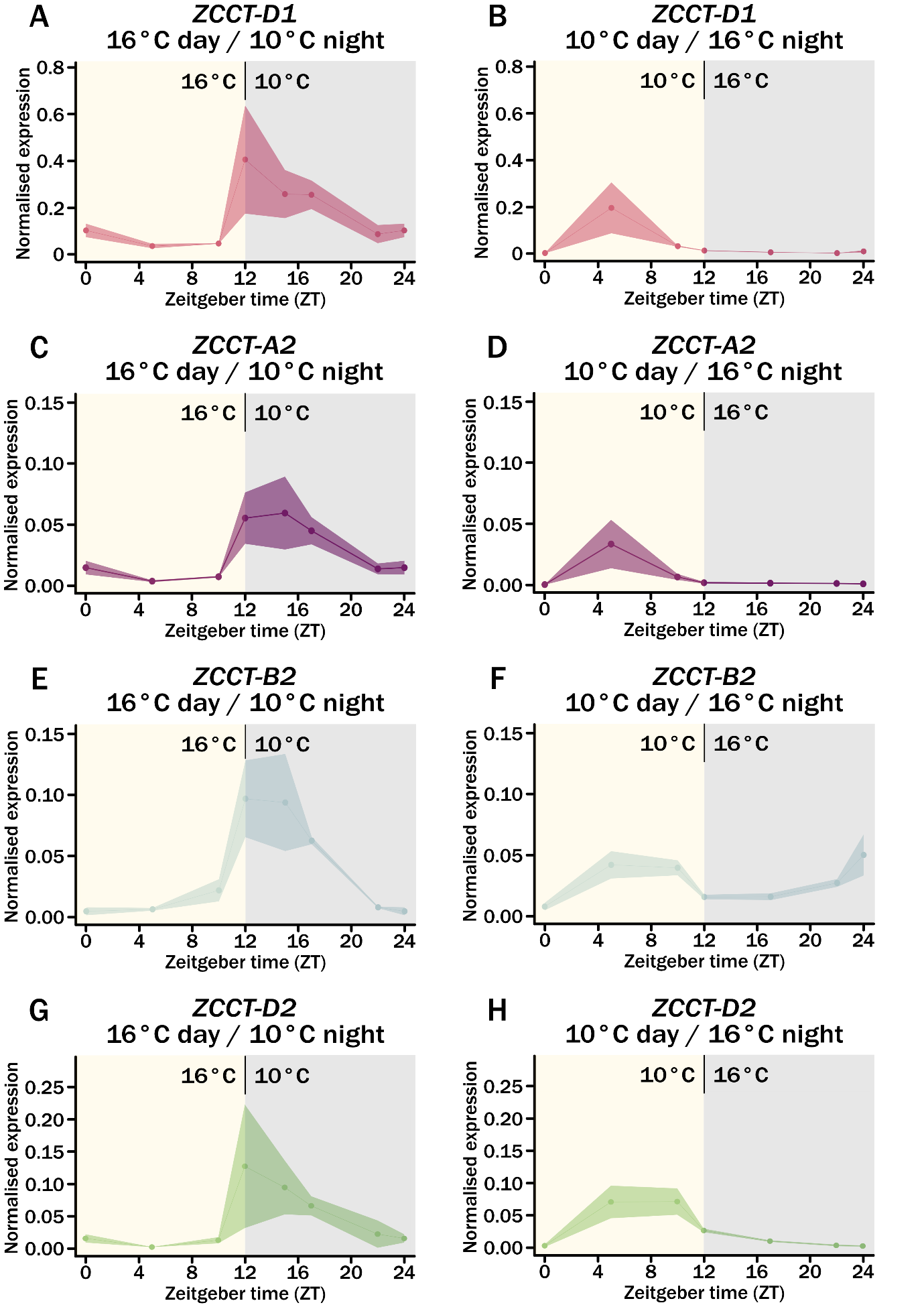

**Figure S2.** Expression pattern of the individual genome copies of *ZCCT1* and *ZCCT2* under varying temperature conditions in a day-neutral photoperiod.

**Table S1.** List of primers

| **Target gene / SNP** | **Technique** | **Genome target** | **Direction** | **Primer sequence (5’ 🡪 3’)** | **Designed by** |
| --- | --- | --- | --- | --- | --- |
| *TraesCS5A02G015600* | RT-qPCR |  | F | TCTAAATGTCCAGGAAGCTGTTA | Borrill et al., 2016 |
|  |  |  | R | CCTGTGGTGCCCAACTATT |  |
| *ZCCT1* | RT-qPCR | All | F | GCAAGAGCCCACATCGTGCC | D. Hirsz |
|  |  |  | R | GCATTGTGGGATAATGGGCAG |  |
| *ZCCT2* | RT-qPCR | All | F | CACCAGCACTATTAGCAATGCAACG | D. Hirsz |
|  |  |  | R | CATTGACCCGTGGCCTGAGCTCG |  |
| *ZCCT-D1* | RT-qPCR | D | F | CCAGCTACGCGAGCACCAGTTCTTC | D. Hirsz |
|  |  |  | R | GTGAGCCTGCTGATGCTGTTCCCTG |  |
| *ZCCT-A2* | RT-qPCR | A | F | AGACGCAGGCGGCCAAAACATG | D. Hirsz |
|  |  |  | R | CATAGCGGATTTGCTTGTCATAGCACC |  |
| *ZCCT-B2* | RT-qPCR | B | F | CCGGCCAATTGCCACCACTGC | D. Hirsz |
|  |  |  | R | CAGCGTCCACCTCCAGGGTC |  |
| *ZCCT-D2* | RT-qPCR | D | F | GACGCGGGCGGACAACACAC | D. Hirsz |
|  |  |  | R | GCCACCATCATCTCTGTATCAATTGTCCTG |  |
| *VRN1* | RT-qPCR | All | F | GAACAAGATCAACCGGCAGGTGAC | Allard et al., 2012 |
|  |  |  | R | GGAGAAGATGATGAGGCCGACCTC |  |
| *ZCCT-D2 Q144* Cadenza0810* | KASP genotyping | D | F | GAAGGTGACCAAGTTCATGCTctgcccataatccgacgatgC | L. Dixon |
|  |  |  | F | GAAGGTCGGAGTCAACGGATTctgcccataatccgacgatgT | L. Dixon |
|  |  |  | R | GTACCTCATCACCTTCGCCTC | L. Dixon |
| *ZCCT-D1 T130I Cadenza1436* | KASP genotyping | D | F | GAAGGTGACCAAGTTCATGCTaggccccaccatcatctttG | L. Dixon |
|  |  |  | F | GAAGGTCGGAGTCAACGGATTaggccccaccatcatctttA | L. Dixon |
|  |  |  | R | TTTACGGAGGTGCATTCACA | L. Dixon |
| *ZCCT-D2 Q144* Cadenza0810* | PCR | D | F | GTACATATCTGTTACCGACAAG | D. Hirsz |
|  |  |  | R | GATCATAGGGCGAAGTTG | D. Hirsz |
| *ZCCT-D1 T130I Cadenza1436* | PCR | D | F | GTGCACCTTTGAATGAAAATGG | D. Hirsz |
|  |  |  | R | GCTGGAGCTCAGCGTAAG | D. Hirsz |

**Table S2.** Weekly growth conditions for long-term gene expression experiment

| **Day** | **Week** | **Temperature (day)** | **Temperature (night)** | **Photoperiod** | **Sample time 1** | **Sample time 2** |
| --- | --- | --- | --- | --- | --- | --- |
| 1 | 1 | 16 | 10 | 13 |  |  |
| 8 | 2 | 16 | 10 | 13 | ZT1 | ZT14 |
| 15 | 3 | 16 | 10 | 13 | ZT1 | ZT14 |
| 22 | 4 | 13 | 8 | 11 | ZT1 | ZT12 |
| 29 | 5 | 9 | 5 | 9 | ZT1 | ZT10 |
| 36 | 6 | 6 | 4 | 8 | ZT1 | ZT9 |
| 43 | 7 | 6 | 4 | 8 | ZT1 | ZT9 |
| 50 | 8 | 6 | 4 | 8 | ZT1 | ZT9 |
| 57 | 9 | 6 | 4 | 8 | ZT1 | ZT9 |
| 64 | 10 | 6 | 4 | 10 | ZT1 | ZT11 |
| 71 | 11 | 8 | 4 | 12 | ZT1 | ZT13 |
| 78 | 12 | 10 | 5 | 14 | ZT1 | ZT15 |
| 85 | 13 | 14 | 7 | 16 | ZT1 | ZT17 |
| 92 | 14 | 17 | 12 | 16 | ZT1 | ZT17 |
| 99 | 15 | 20 | 14 | 16 | ZT1 | ZT17 |
| 106 | 16 | 20 | 14 | 16 | ZT1 | ZT17 |
| 113 | 17 | 20 | 14 | 16 | ZT1 | ZT17 |
| 120 | 18 | 20 | 14 | 16 | ZT1 | ZT17 |
| 127 | 19 | 20 | 14 | 16 |  |  |
| 134 | 20 | 20 | 14 | 16 |  |  |

**Table S3.** List of germplasm

| **Genotype** | **Growth habit** | **Country of origin** | **Background/ Pedigree** | **Source** |
| --- | --- | --- | --- | --- |
| Cadenza | Facultative spring | UK | Axona / Tonic | Dixon lab, University of Leeds |
| Cadenza0810 BC2F2 |  | UK | Cadenza x [Cadenza x Cad0810] | Dixon lab, University of Leeds |
| Cadenza1436 BC2F2 |  | UK | Cadenza x [Cadenza x Cad1436] | Dixon lab, University of Leeds |
